# Self-supervised image restoration in coherent X-ray neuronal microscopy

**DOI:** 10.1101/2025.02.10.633538

**Authors:** Alfred Laugros, Peter Cloetens, Carles Bosch, Richard Schoonhoven, Liam Pavlovic, Aaron T. Kuan, Jayde Livingstone, Yuxin Zhang, Minsu Kim, Allard Hendriksen, Mirko Holler, Adrian A. Wanner, Anthony Azevedo, K. Joost Batenburg, John C. Tuthill, Wei-Chung Allen Lee, Andreas T. Schaefer, Nicola Vigano, Alexandra Pacureanu

## Abstract

Coherent X-ray microscopy is emerging as a transformative technology for neuronal imaging, with the potential to offer a scalable solution for reconstruction of neural circuits in millimeter sized tissue volumes. Specifically, X-ray holographic nanotomography (XNH) brings together outstanding capabilities in terms of contrast, spatial resolution and data acquisition speed. While recent XNH developments already enabled generating valuable datasets for neurosciences, a major challenge for reconstruction of neural circuits remained overcoming resolving power limits to distinguish smaller neurites and synapses in the reconstructed volumes. Here we present a self-supervised image restoration approach that simultaneously improves spatial resolution, contrast, and data acquisition speed. This enables revealing synapses with XNH, marking a major milestone in the quest for generating connectomes of full mammalian brains. We demonstrate that this method is effective for various types of neuronal tissues and acquisition schemes. We propose a scalable implementation compatible with multi-terabyte image volumes. Altogether, this work brings large-scale X-ray nanotomography to a new precision level.

## 1. INTRODUCTION

In the quest for fundamental understanding of brain function, for solutions to neurological disorders, or for optimal design of neuromorphic chips, scientists coordinate efforts into pushing technology to generate neuronal wiring diagrams [1]. While electron microscopy is the established imaging method for generating image volumes with sufficient precision to reconstruct neuronal circuits [2], probing brain tissue with hard X-rays offers several benefits. These include penetration through multi-millimeter sized tissue thickness, preservation of the whole sample post-imaging, and wavelength of the illumination in the sub-nanometer range. By using a coherent beam, hard X-ray phase contrast imaging offers orders of magnitude improvement in contrast in soft tissues [3]. The advent of fourth generation synchrotron sources makes available highly coherent X-ray probes with photon flux enabling substantial acceleration of data acquisition rates [4]. Leveraging these advances, X-ray holographic nanotomography (XNH) [5, 6] emerges as a suitable technology for imaging brain tissue with nanometer resolution at large scale. While valuable neuronal tissue data could be recently generated with XNH [7–11], the spatial resolution in brain tissues remained limited to around 80 nm. These XNH data sets helped us to understand how the nervous system controls the body ([12, 13]) and to bridge the gap between the brain microstructure and functional magnetic resonance measurements that can be conducted safely in patients ([14]). X-ray ptychographic nanotomography [15, 16], also a coherent X-ray microscopy technique, was also adapted for neural tissue imaging [17, 18], and was recently demonstrated to achieve sub-40 nm resolution [19]. For X-ray microscopy, the interest of XNH in the context of connectomics is a gain in data collection speed of several orders of magnitude. Here we present an approach for self-supervised image restoration that makes it possible to attain an important milestone in terms of resolving power of scalable X-ray nanotomography. Convolutional neural networks have demonstrated remarkable efficacy in denoising images [20]. They can learn to remove various types of artifacts using pairs of ground truth and corrupted images [21]. However, obtaining ground truth data can be challenging, particularly in the field of biological imaging. To address this issue, methods that do not require paired clean data have been developed. For example, content-aware image restoration (CARE) [22] approaches leverage unpaired high signal-to-noise ratio (SNR) images to denoise those with lower SNR. Nevertheless, obtaining high SNR data is not always feasible. Moving forward, methods that can restore images based on only noisy data, such as Noise2Void [23] and Noise2Self [24] have been proposed. However, these approaches require that the noise to be removed is element-wise independent, an assumption that is incompatible with XNH data, where reconstructed image volumes contain structured noise.

More recent self-supervised approaches have been developed to remove as well pixel-wise dependent noise [25–27]. These methods remain however not entirely noise-agnostic. They require tuning of various parameters, and the restoration performance for each set of parameters is strongly dependent of the characteristics of the noise to be removed. The alternative approaches for removing structured noise are based on pairs of noisy images, such as Noise2Noise (N2N) [28]. In the context of X-ray tomography, Noise2Inverse [29] has been developed and applied to restore images in the reconstruction domain [30, 31]. Here, we build up on this concept, and develop a 3D scalable approach for restoring XNH image volumes of neuronal tissue. This development enables identifying synapses with X-ray holographic nanotomography, a technology that is already capable of generating valuable data sets in millimeter-sized samples. The machine learning based advances we report here offer simultaneous gains in spatial resolution, contrast, and data collection speed, without requiring high-quality ground truth data. In multi-terabyte image volumes, transfer learning enables substantial acceleration of computation time and resource savings. Thanks to a full 3D implementation, structured noise resulting from mixing of the illumination probe and the imaged object is effectively removed. Importantly, we demonstrate that ring artifacts that are persistent in X-ray tomography and have remained a long-standing problem, are effectively removed.

## 2. RESULTS

### A. Restoration Performance

The self-supervised restoration approach we propose here is suitable for 3D X-ray nanoimaging, and more specifically for X-ray holographic nanotomography (XNH) (Supplementary Figure 1). Conceptually, the method is based on the Noise2Noise (N2N) [28] principle adapted to X-ray tomography [29]. We developed a framework that offers substantially improved performance both in terms of restoration precision and scalability. To quantify the performance of the restoration or denoising, terms that we use interchangeably, we measured the Contrast to Noise Ratio (CNR) [32] and the spatial resolution as estimated with Fourier Shell Correlation (FSC) [33]. Further details about these metrics are provided in Section E. Table 1 summarizes these measurements before and after denoising image volumes of mouse and fruit fly neuronal tissue. Additional information on the specimens and XNH data generation is included in section B). We obtain contrast to noise ratio enhancement of up to five fold, and resolution improvement of more than a factor two for the fly tissue sample.

**Fig. 1.**
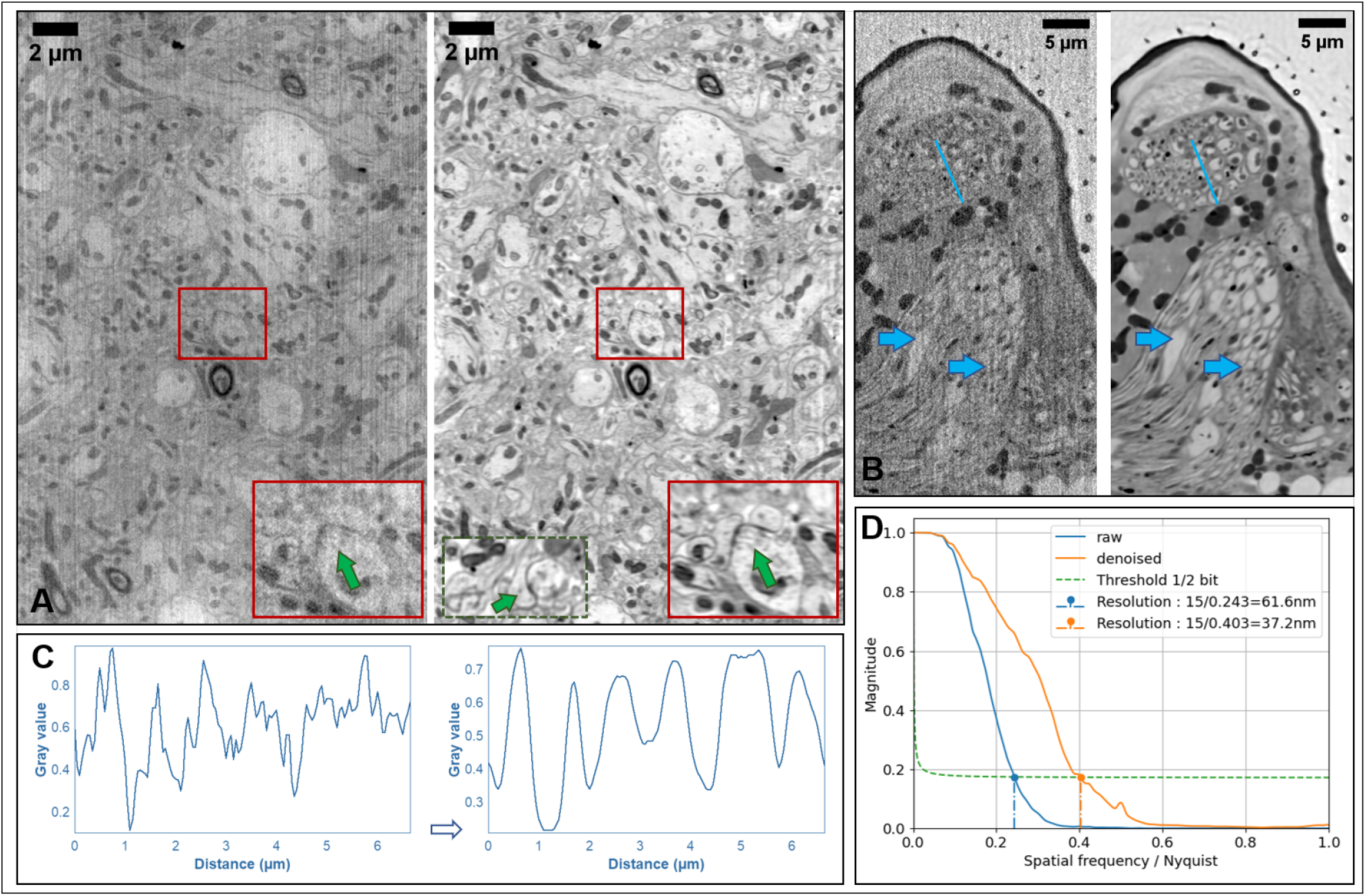
Illustration of the performance of the proposed approach on mouse and fruit fly neuronal tissue. A. Matched reconstructed sections from a mouse olfactory bulb tissue sample (OB15), before and after image restoration. The magnified sub-region marked with a rectangle includes two synapses (green arrow). The dotted border inset shows an orthogonal view through one of the two synapses present in the magnified sub-region. B. Restoration of neuropil in fruit fly tissue (FSS). The displayed region includes the fly halter nerve. Arrows point to areas affected by structured noise, before and after restoration. C. Line profiles across the regions marked in panel B, showing that after denoising, a series of consecutive neurites become well distinguishable. D. Fourier shell correlation plots showing the improvement in measured spatial resolution for the data corresponding to panel (A).

The resolution and contrast gains make is possible to identify synapses (Figure 1 A). The magnified regions show two synapses, together with an orthogonal view through one of them. Since XNH data offers isotropic spatial resolution, the image quality is preserved along the three orthogonal directions. Note that the extra-cellular space is resolved between the densely packed neurites. The resolution improvement is also reflected in the FSC curves, as it can be seen in Figure 1 D. The neurites in the fly halter nerve shown in Figure 1 B become distinguishable after denoising. This is confirmed by the clearly separated peaks in the gray level line profiles shown in panel C.

Standing out from other approaches, the method proposed here both separates and removes structured noise from the 3D images. This can be observed in Figure 1, panels A and B, with blue arrows indicating some of the stripe type of artifacts that are effectively removed without compromising the useful signal. In XNH, these residual stripes arise from the mixing of the X-ray probe with the imaged object [34]. The ability of this self-supervised approach to unmix the probe-object signal is of particular importance for XNH data reconstruction.

Thanks to the image quality enhancement, multiple biological features become well resolved. Inside the cell body, as seen in Figure 2 A containing a neuron from the external plexiform layer (EPL) tissue, we can identify organelles such as the endoplasmic reticulum, mitochondria, and the nuclear membrane. These structures are also visible in the volume rendering displayed in 2 B. A segmented neurite together with a synapse are displayed in panel C.

**Fig. 2.**
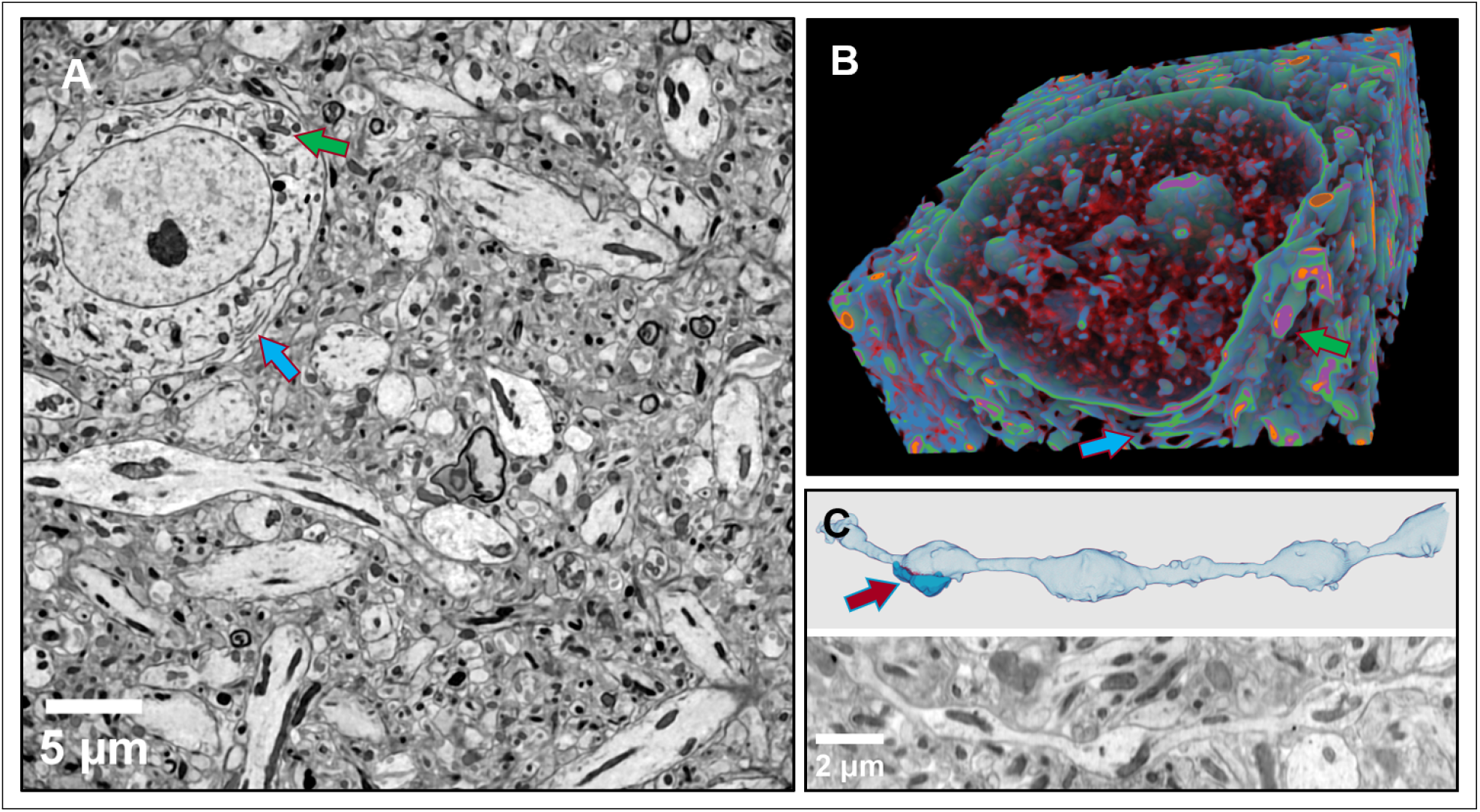
A. Section from a restored 3D image of a mouse olfactory bulb sample (OB25), including a neuron cell body. Organelles such as endoplasmic reticulum (ER) (blue arrow) and mitochondria (green arrow) are clearly distinguished. B. Volume rendering of a region corresponding to the cell body seen in panel (A) showing detailed structures such as chromatin, nuclear membrane, ER (blue arrow) and mitochondria (green arrow). C. Volume rendering of a segmented neurite containing several synaptic hubs (red arrow) together with a corresponding section through the image volume.

An important contribution of the approach we propose here is the effective suppression of ring artifacts, a long standing problem in X-ray tomography [35, 36]. Figure 3 illustrates the ring removal results obtained on a mouse hippocampus tissue sample. Orthogonal sections through the image volume before and after restoration show the ability of the proposed method to untangle and eliminate this structured 3D noise without altering the features of the imaged tissue. Although we demonstrate it here for XNH data, the method is readily applicable to any type of X-ray tomography approach. A comparison of the results obtained with state-of-the-art methods and our approach is included in Supplementary Figures 4 and 5. The improvement in the performance is prominent.

**Fig. 3.**
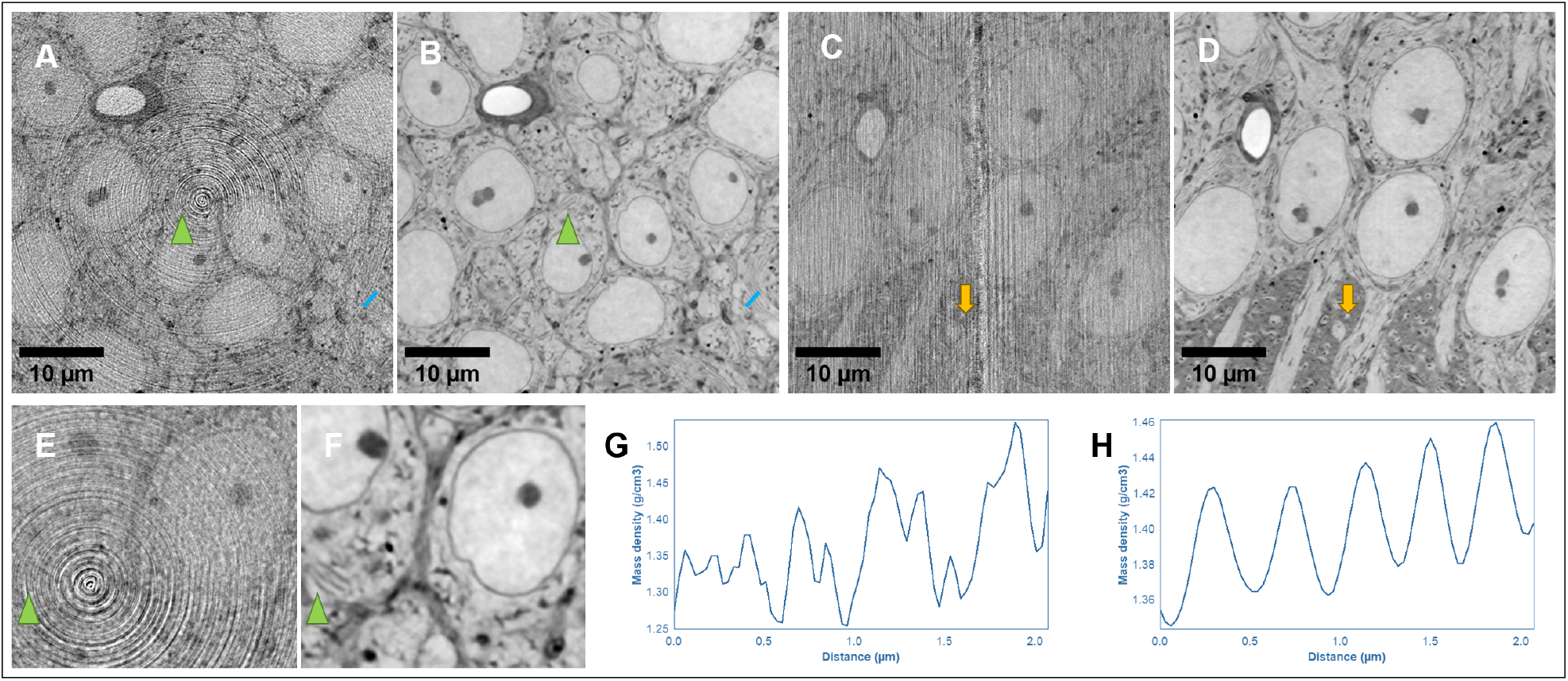
Illustration of the removal of structured ring noise, an artifact inherent in tomography. A,B and C,D. Paired orthogonal slices before and after self-supervised denoising of images from mouse hippocampus (HPO). E and F. Detailed regions form panels A and B, respectively. G and H. Line plots along the region marked over panels A and B, highlighting the restoration of endoplasmic reticulum inside a neuron. Other sub-cellular structures, such as nuclear membranes and mitochondria emerge clearly once the noise is removed. Matched arrows and arrowheads point to paired structures before and after restoration.

### B. Training Time and Scalability

Table 1 shows the training times of models using two Nvidia A100 GPUs. We note that training time scales with training volume size: the HPO volume is fifty times larger than the FSS volume, yet the training on HPO is only five times longer than the training on FSS. Additional results demonstrating the scalability of our denoising implementation are provided in the Supplement 4.3.

**Table 1.** Denoising performances

| Volume | Measured Resolution | CNR | Training Time |
| --- | --- | --- | --- |
| Raw FSS | 235.8 | $1.9 \pm 0.4$ | - |
| Denoised FSS | 96.1 | $9.8 \pm 1.2$ | 4h |
| Raw OB15 | 61.7 | $2.8 \pm 0.2$ | - |
| Denoised OB15 | 37.2 | $7.8 \pm 1.1$ | 13h |
| Raw OB25 | 91.9 | $3.5 \pm 0.5$ | - |
| Denoised OB25 | 51.0 | $7.5 \pm 0.4$ | 14h |
| Raw HPO | 196.1 | $1.4 \pm 0.4$ | - |
| Denoised HPO | 103.8 | $6.8 \pm 0.9$ | 20h |

The values in Table 1 correspond to the training duration required to achieve the best denoising performances, but trainings already achieve good results long before convergence. For instance, the best resolution we measure for the FSS sample is 140 nm, and it is obtained after four hours training. But after only one hour training, we already measure a resolution gain of 100 nm. In Supplementary Figure 6, we display the resolution measured at various stages of trainings for the FSS and OB15 samples. We measure that resolution improves much faster at the beginning of the trainings than at the end. Therefore, to save computational resources, training can be stopped long before convergence by accepting a small loss in terms of denoising performances.

Training time can also be reduced by using a pre-trained model that can be rapidly fine-tuned to restore a new image volume. Specifically, the time required to fine-tune a model that has been trained on one volume from a sample to another volume from the same sample, is ten times faster than a training from scratch. The obtained denoising performances remain unaltered.

### C. Data acquisition strategies

The N2N algorithm requires two statistically independent observations of the signal to be denoised [28]. Hendriksen et al. propose obtaining these data by partitioning the angular projections acquired during tomographic scanning into different subsets [29], with each subset used independently to reconstruct a volume. We used this strategy to obtain training volumes for the FSS, OB15 and OB25 data sets. In addition, we propose two new approaches to generate independent observations of the sample.

The first approach consists in acquiring two consecutive continuous, or on-the-fly tomographic scans with a slight sample position shift in-between (Supplementary Figure 1). This results in two image volumes of essentially the same region of the specimen, affected by statistically independent noise. For each of the two scans, the exposure time per frame is reduced by 50% to maintain the same total radiation dose. Since during continuous acquisition mode the rotation motor is never stopped during the angular data collection, the overheads related to moving actuators are substantially reduced. This results in an effective gain in data collection time of around a factor two. The reconstructed volumes are, however, affected by ring artifacts that are inherent to X-ray tomography [36]. For step-by-step tomographic scanning, a lateral sample displacement at each angular position offers a solution [34], at the price of longer acquisition duration. Here we propose a single sample lateral displacement between the two tomographic scans, resulting in a shifted location of the ring artifacts relative to the specimen features. An detailed illustration is provided in Supplementary Figure 8. This strategy offers an effective solution for removing ring artifacts, as seen in Figure 3). The HPO sample has been acquired with this approach and we observe in Table 1 that denoising performances obtained on this volume are similar to those achieved when using the angular projection partitioning.

The second method we propose to generate independent representations of the useful signal is based on the longitudinal diversity for phase retrieval. For XNH, we typically acquire projections at four propagation distances [6, 7] (Supplementary Figure 1). These four holograms are aligned and combined together to obtain phase maps [5]. We propose to use two pairs of projections recorded at different propagation distances to generate two phase maps for each angular position. This gives rise to two reconstructed volumes that are based on data acquired at the same angular positions, but from different propagation distances. As shown in Figure 10 and Table 5 of the Supplement, the denoising performances of models trained using longitudinal phase diversity are comparable to those achieved using the angular projection partitioning strategy. This approach does not require any change in data collection strategy, and it is applicable as well to the continuous scanning method presented above. Supplementary Table 1 includes details about the acquisition settings.

## 3. DISCUSSION

Holographic X-ray nanotomography is becoming a transformative technology for neuronal imaging. The self-supervised image restoration approach that we propose here improves the spatial resolution by up to a factor two, and the contrast by up to a factor five. These improvements make possible the identification of synapses with XNH, paving the way towards reconstruction of neuronal circuits in multi-millimeter sized tissue volumes with X-ray microscopy. Exceptional efforts and advances in developing volume electron microscopy enabled recently the creation of a few extraordinary data sets [1, 37–39]. The merits of X-ray imaging include the ability to work with large tissue samples [40–42] circumventing the need for ultra thin serial sectioning or ablation, while offering the potential to reach sub-ten nanometers spatial resolution [43]. Combined with the inherent alignment and isotropic resolution of the generated volumes, this can greatly accelerate data collection speed for connectomics and for nanoscale tissue investigation more broadly.

The results presented here push the limits of achievable precision for imaging neuronal tissue at nanometer scale in samples with thickness in the range of hundreds of micrometers. A notable contribution is the ability to untangle structured noise, such as residual mixing of the X-ray nanoprobe with the specimen structures. The strategy we propose to remove ring artifacts is directly applicable to X-ray tomography in general, and thus of interest to a large community. The developments described here make it possible to work with reduced radiation dose and to accelerate data acquisition. For equivalent measured spatial resolution in the range of 40 nm, as reported in [19], we could reduce the radiation dose to one third, and cover a volume more than two orders of magnitude larger during the same acquisition time. Several avenues can be envisaged to further improve spatial resolution for neural imaging with XNH. A fist computational avenue is addressing the residual sample deformation that occurs during data collection [44]. A second perspective is including the illumination probe retrieval within the XNH phase reconstruction process [45–47]. For scaling up data acquisition, complementary advances in instrumentation such as photon-counting X-ray detectors, and high precision actuators, can play a major role.

In summary, the advances presented here push the spatial resolution frontier in XNH, offer a robust tool for improving data quality in X-ray tomography more broadly, and lay the foundation for large scale X-ray neuronal imaging, with particular impact in connectomics.

## 4. METHODS AND MATERIALS

### A. Neuronal tissue samples

The tissue samples included here are from mouse olfactory bulb (OB), mouse hippocampus (HPO), and *Drosophila* flight stabilization system (FSS). All tissues were prepared with protocols tailored for volume Electron Microscopy, including fixation, metal staining and resin embedding. A detailed description is available in [7, 48]. The OB samples were cut into pillars using either the system described in [49] or a focused ion beam. The HPO sample was cut into a rectangular prism using a vibratome. The FSS sample encompassed the fly haltere, the haltere nerve, and the ventral nerve cord, and had the native shape of the anatomical structures.

### B. X-ray nanotomography data acquisition and reconstruction

We acquired the X-ray nanotomography data at the ID16A beamline of the ESRF [50, 51]. The main acquisition parameters are summarized in Table 2. For sample OB25 the detector was composed of a 23-micron thick GGG-Eu scintillator and a Frelon CCD camera. For the remaining samples included here, the detector was composed of a 20 microns thick LSO-Tb scintillator coupled to a Ximea sCMOS camera. To obtain each three dimensional image, four sets of angular projections over 180 degrees were recorded at different propagation distances, as illustrated in Supplementary Figure 1. To obtain phase maps, the four longitudinally diverse holograms corresponding to each rotation angle were aligned and combined using an iterative algorithm based on conjugate gradient descent [7]. The final set of phase maps was used for tomographic reconstruction based on filtered backprojection.

**Table 2.** Data acquisition parameters

| Dataset | Energy [keV] | Voxel size [nm] | Angular projections | Exposure time [s] | Dose [Gy] |
| --- | --- | --- | --- | --- | --- |
| OB25 | 33.6 | 25 | 1800 | 0.15 | $8.4 \times 10^7$ |
| OB15 | 33.6 | 15 | 3000 | 0.15 | $8.5 \times 10^8$ |
| FSS | 33.6 | 50 | 2000 | 0.25 | $8.1 \times 10^7$ |
| HPO | 33.6 | 30 | 3000 | 0.30 | $3.3 \times 10^8$ |

### C. Training Process

For training the neural network, we selected subvolumes that contain mostly tissue, leaving aside empty resin regions that are contained within the reconstructed 3D images. To obtain the image volume pairs used as input, we applied one of the three approaches described in Section C: angular projection partitioning, propagation distance partitioning, or shifted tomographic acquisitions. More details regarding the training volumes are included in the Supplementary Table 1. From these volumes, we extracted pairs of randomly selected 96^3^ voxels patches. For each training pair, a random orientation was selected from the 24 possible rotational symmetries of a cube. The patches were then rotated accordingly. Finally, a horizontal flip was applied with a 50% probability.

The trained models were 3D U-Nets [52] with four convolution blocks in the encoder pathway. The initial convolutional layer had 56 filters. We used instance normalization instead of batch normalization, as we found that this leads to better denoising performance.

The networks were trained using L2 loss and the Adam optimizer. The learning rate was set to 5∗ 10^*−*4^ and the batch size to 32. Models were trained until convergence, that is until denoising performance stops improving. The hyper-parameter values were determined through successive random searches. Additional details are provided in the Supplementary material section 5.

The implementation we propose here generates substantially improved results compared to the Noise2Inverse approach based on the Mixed Scale Dense network architecture [53]. Quantitative results are included in Supplementary section 4.

### D. Inference Process

Onced trained, models are applied to volumes reconstructed using all projections as recommended in [54]. However, when using the continuous scan acquisition strategy (see Section C), there are no projection splits to be combined since training volumes are acquired with two distinct scans. Therefore, in this case, we use the original Noise2Inverse inference procedure [29], i.e. the two training volumes are processed separately, and the mean of the processed volumes constitutes the final result.

Volumes to be denoised are processed patch-by-patch. Processed patches are aggregated to reconstruct denoised volumes. For each dimension, the stride used to extract successive patches equals 80% of the training patch size. This means that extracted patches overlap. We use a stride smaller than the patch size to prevent artifacts at the borders of aggregated patches (examples are displayed in Supplementary Figure 9). A Hann window function is used to weight patches in overlapping regions.

### E. Performance Evaluation Process

To evaluate image restoration or denoising performance, we used the three criteria described below.

1. *Contrast to Noise Ratio (CNR)* metric [32] defined as:

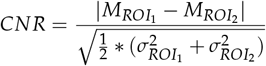

where 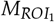 and 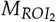 are the mean gray values of the regions (*ROI*_1_ and *ROI*_2_) that we aim to distinguish in the image volumes. We manually selected *ROI*_1_ and *ROI*_2_ across various sections of each volume (an illustration is provided in Supplementary Figure 7). We then calculated the CNR for each section and took the average to determine the overall CNR of the volume. The confidence intervals of the CNR of volumes are given with a confidence coefficient of 95%. The two types of regions of interest we selected for these measurements are cell membranes and cell cytosol.
2. *Measured Resolution* using Fourier Shell Correlation (FSC) [55]. We applied the neural networks to the two image volumes used for training. We then used the obtained images to compute the FSC curves. The measured resolution was determined using the *1/2-bit threshold* criterion [33]. The obtained FSC curves before and after image restoration for volume OB15 are shown in Figure 1 D. The evolution of the FSC measured resolutions as function of training time is illustrated in Supplementary Figure 6.
3. *Visual evaluation*: Based on human expertise, we assessed the improvement in visual quality of the biological structures we aim to investigate.
4. The code used to train and evaluate deep neural networks in our experiments can be found here : https://github.com/xniesrf/SSD_3D.

### F. Additional software

For tomographic reconstructions, including ring artifact correction using methods from literature, we used the Nabu software (https://gitlab.esrf.fr/tomotools/nabu). For phase retrieval we used GNU Octave [56]. For 3D image alignment prior to ring correction we used the pi2 library (https://github.com/arttumiettinen/pi2) [57]. For figure preparation and for selection of image volume regions for CNR measurements we used Fiji [58]. For volume renderings appearing in Figure 2 we used Paraview (https://www.paraview.org/). For semi-automatic image segmentation we used ilastik [59].

## Acknowledgments

This work was supported by the European Commission through the ERC project BRILLIANCE (852455) to AP. We acknowledge funding from UKRI Physics of Life initiative (EP/W024292/1) to AP and ATS and from NIH BICCN (MH128949) to AP and WCAL. We are grateful to Pierre Paleo for implementing into Nabu ring artifact removal methods from literature. CB, YZ and ATS are supported by the Francis Crick Institute, which receives its core funding from Cancer Research UK (FC001153), the UK Medical Research Council (FC001153), and the Wellcome Trust (FC001153). For the purpose of Open Access, the author has applied a CC BY public copyright licence to any Author Accepted Manuscript version arising from this submission. We are grateful to the European Synchrotron for awarding beamtine for proposals LS2892, LS3230 and LS3231.

## Author contributions

Conceptual design: AL, PC, NV, AP. Implementation: AL with contributions from RS, LP, AH, NV. Sample preparation: CB, ATK, MK, MH, AAW. Data acquisition: PC, CB, JL, AA, ATS, AP. Data processing and analysis: AL, PC, JL, AP. Funding acquisition and supervision: JB, JCT, WCAL, ATS, AP. Manuscript preparation: AL and AP with contributions from all authors.

## Data Availability Statement

The code with data are available at https://github.com/xni-esrf/SSD_3D.

## Supplementary document

See the Supplementary file for supporting content.

