## Supplementary file for "Self-supervised image restoration in coherent X-ray neuronal microscopy"

### Supplementary material

#### 1. The acquisition setups

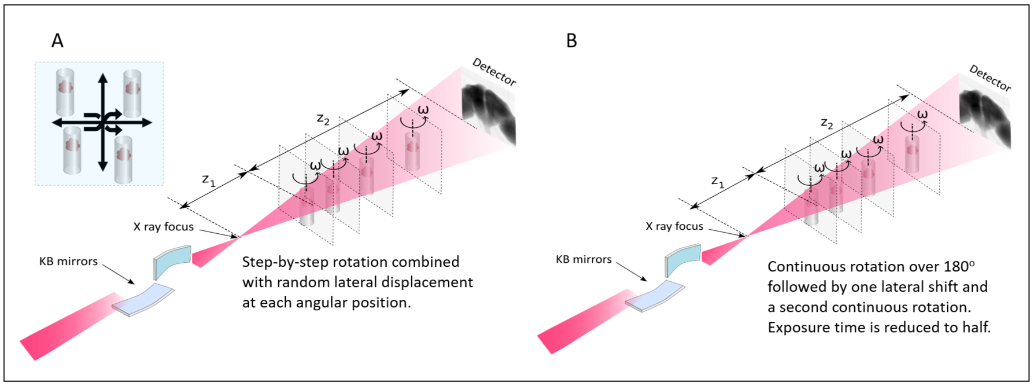


Fig. 1. Schematics of the two experimental setups used for data XNH acquisition.

The first acquisition strategy relies on step-by-step tomographic acquisition combined with random lateral displacement (panel A). In this case the rotation is paused for each frame and nanopositioning piezo actuators are used to move the sample orthogonal to the incident beam by a few pixels ( [1,2]). The second strategy is based on continuous tomographic acquisition with two full consecutive rotations (panel B). In this case the rotation is not interrupted during frame acquisitions. The sample is slightly shifted in-between the two tomographic scans to obtain a different relative position of the structured artifacts arising from the mixing of the probe and the object. The exposure time for the two consecutive continuous acquisitions is reduced to half compared to the step-by-step scans, in order to conserve the total radiation dose.

#### 2. Details of the volumes used for training

Table 1. **Partitioning strategy and crop size of training volumes**

| Training volumes | Partitioning Strategy | Crop Size |
|---|---|---|
| OB15 | Angle | 1800x1200x1200 |
| FSS | Angle | 500x500x500 |
| HPO | Two Shifted Scans | 1900x1800x1800 |
| OB25 | Angle | 1500x1500x1500 |
| OB25-dist | Distance | 1500x1500x1500 |

#### 3. Supplementary examples of synapses

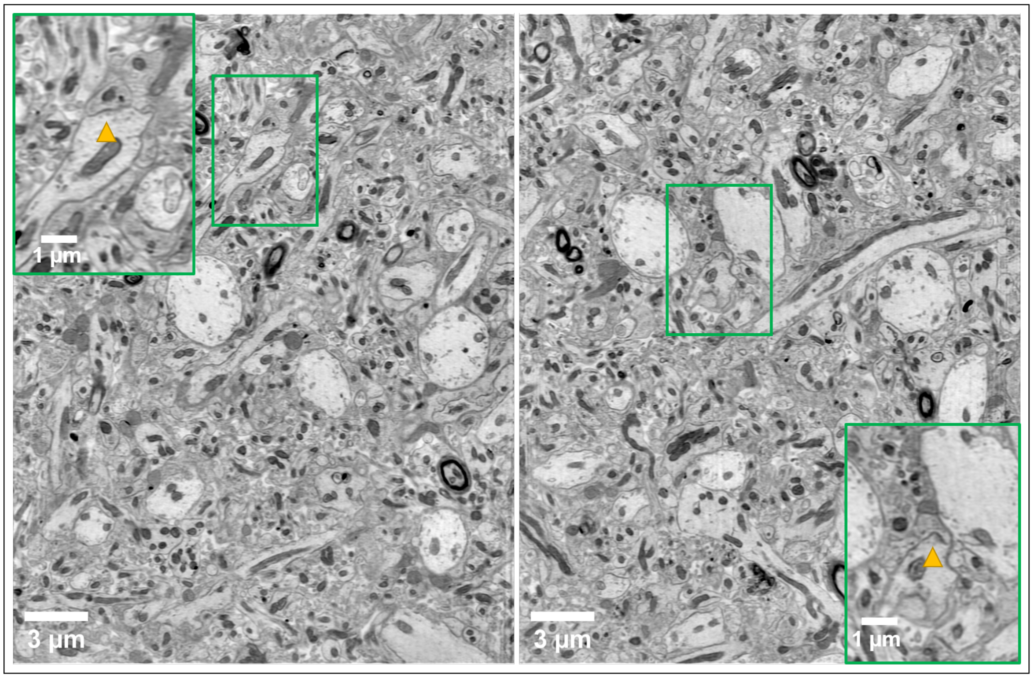


Fig. 2. Additional examples of synapses indicated with arrows in the magnified regions.

#### 4. Comparison between 3D U-Net and MSD implementations

In this Section we compare our implementation based on the 3D U-Net architecture described in Section 4, with a previous implementation based on the Mixed Scale Dense (MSD) network [3]. To achieve that, we consider two volumes: FSS and OB25 described in Section B. In this Section, the volumes required to train models are obtained using the standard Noise2Inverse angle split procedure [4].

##### *4.1. The MSD training set-up*

The MSD networks trained in our experiments have the same structure and number of parameters as described in [4]. Note that MSD processes volumes slice by slice, meaning the data loading pipeline used with MSD differs from that used with 3D U-Nets (which are trained using 3D patches).

Each pair used to train MSD networks is obtained by randomly picking a location, and then by extracting from the two training volumes the slice having this location. We use some data augmentation to increase training data diversity. Slices are extracted from three directions of the training volumes with equal probability, rotated by 0°, 90°, 180°, or 270° with equal probability, and horizontally flipped with a probability of 0.5. Data augmentation significantly improves the performance of the trained MSD.

We also apply a technique presented in [5] that has been shown to increase denoising performances. This technique consists in joining context slices to input slices for each input-target training couple. This formatting enables trained networks to better leverage information from the

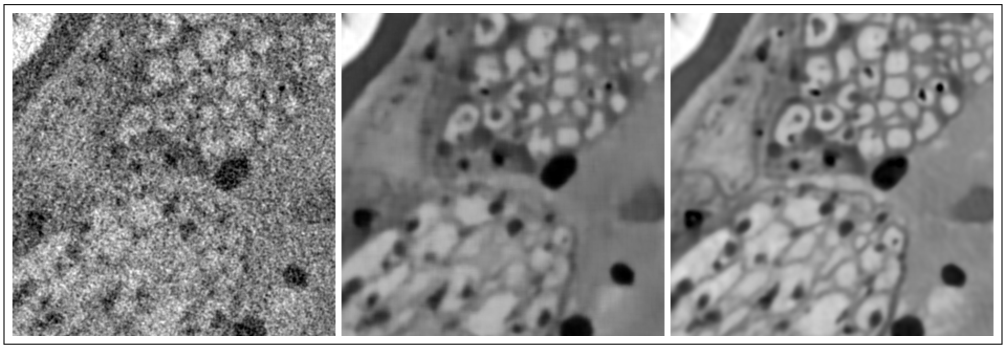

Fig. 3. Crops of the FSS volume. Left: raw data. Middle: denoised using the MSD implementation. Right: denoised using the 3D U-Net implementation.

dimension orthogonal to input slices. In our experiments, we add four context slices to every input slice.

MSD networks are trained using the L2 loss function, the Adam optimizer with a learning rate of 0.001, and a batch size of 16. During inference, volumes denoised by MSD are processed slice by slice, unlike the patch-by-patch procedure used for 3D U-Net described in Section D).

##### 4.2. Denoising Performances

The denoising performances of the MSD and the 3D U-Net implementation are compared in Table 2. For both the FSS and OB25 volumes, the CNR and resolution measured are significantly better with the 3D U-Net implementation.

When examining the denoised volumes, it is clear that better results are achieved using the 3D U-Net implementation (see Figure 3). In particular, fine-grained details are better retrieved with the 3D U-Net implementation compared to the MSD one.

Table 2. Comparison between the denoising performances obtained using the MSD implementation and the ones got using the 3D U-Net implementation.

| Volume | Measured Resolution | CNR | Training Time |
|---|---|---|---|
| Raw FSS | 235.8 | 1.9 ± 0.4 | - |
| FSS denoised using the MSD implementation | 179.2 | 6.0 ± 1.0 | 306min |
| FSS denoised using the 3D U-Net implementation | 96.1 | 9.8 ± 1.2 | 224min |
| Raw OB25 | 91.9 | 3.5 ± 0.5 | - |
| OB25 denoised using the MSD implementation | 55.4 | 6.4 ± 0.2 | 88h |
| OB25 denoised using the 3D U-Net implementation | 51.0 | 7.5 ± 0.4 | 14h |

##### 4.3. Scalability

Training times for the MSD and U-Net models on the FSS and OB25 volumes are displayed in Table 2. For the FSS sample, training time is 30% shorter with the 3D U-Net implementation. The training time difference is even greater for the OB25 sample, suggesting that the 3D U-Net implementation scales better with volume size.

To verify this, we train a MSD model and a 3D U-Net model on three volumes of different sizes: the $500^3$ central crop of the OB25 volume, the $1000^3$ central crop of the OB25 volume and the entire OB25 volume ($1500^3$). The training times for both implementations on these volumes are shown in Table 3. We note that the training time difference between the two implementations increases exponentially with volume size, which confirming that the 3D U-Net implementation scales much better than the MSD implementation.

Table 3. Training times of the MSD and 3D U-Net implementations for several OB25 crop sizes.

| Training Setting \ OB25 Crop Size | $500^3$ | $1000^3$ | $1500^3$ |
|---|---|---|---|
| MSD | 5h | 24h | 88h |
| 3D U-Net | 3h | 8h | 14h |

#### 5. Setting training hyper-parameters with successive random searches

To determine the hyper-parameters for our 3D implementation we conduct three successive random searches. The considered hyper-parameters are: the learning rate, the batch size, the patch size, the number of filters in the first convolution of trained 3D U-Nets, and the number of convolution blocks in 3D U-Nets. For each random search, each hyper-parameter is associated with a set of values. These initial sets of values have been established based on successful implementations of 3D U-Nets in the field of biological imaging.

We test fifteen settings in each random search. A setting is obtained by randomly picking a value for each hyper-parameter. A model is trained for each picked setting, and its denoising performances are measured. The measured denoising performances are used to refine the hyper-parameter ranges to be tested in the next random search. The idea is to increase the probability of sampling the best performing settings. The sets of hyper-parameter values used for the three conducted random searches are provided in Table 4. Our final hyper-parameter setting corresponds to the best performing training of the third random search. It is the setting given in Section C. In these random searches, models are trained on the OB25 volumes.

Table 4. Value sets considered in the conducted random searches for every hyper-parameter

| Hyper-Parameter to set | Random search1 | Random search2 | Random search3 |
|---|---|---|---|
| First convolution filter number | $\{8, 16, 32\}$ | $\{32, 40, 48\}$ | $\{48, 56, 64\}$ |
| Convolution block number | $\{3, 4, 5, 6, 7\}$ | $\{3, 4, 5\}$ | $\{3, 4\}$ |
| Batch Size | $\{4, 16, 64, 256\}$ | $\{4, 16, 64\}$ | $\{8, 16, 32\}$ |
| Learning Rate | $\{10^{-4}, 10^{-3}, 10^{-2}\}$ | $\{10^{-4}, 5 * 10^{-4}, 10^{-3}\}$ | $\{5 * 10^{-4}\}$ |
| Patch Size | $\{96, 128, 160, 192\}$ | $\{64, 96, 128, 160\}$ | $\{64, 96\}$ |

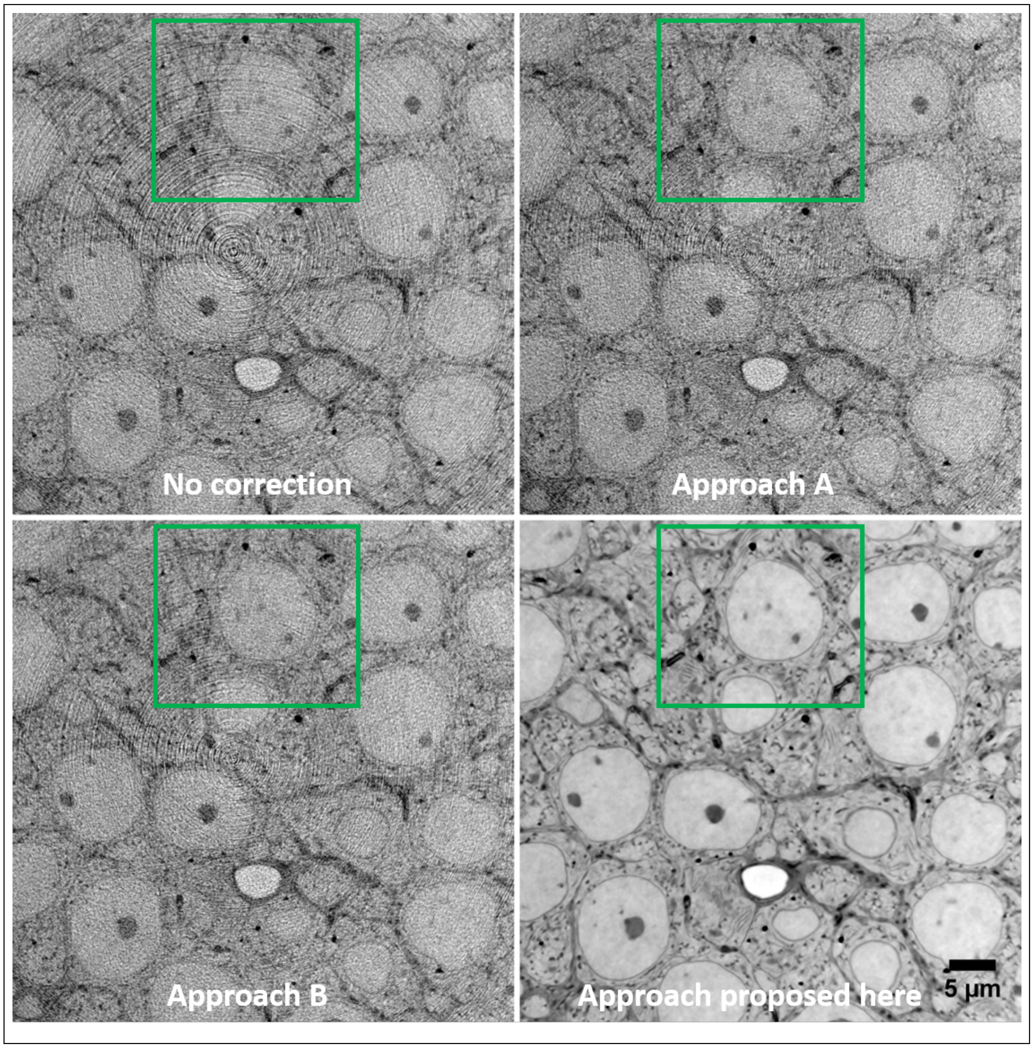


Fig. 4. Results obtained with different ring artifact removal approaches illustrated on a slice from the HPO sample. Top left: no correction. Top right: correction obtained with the method presented in Munch et al. [6]
. Bottom left: correction obtained with the method presented in Vo et al. [7]. Bottom right: self-supervised correction approach proposed here. The two methods existing in the literature are implemented in the Nabu tomographic reconstruction software (https://gitlab.esrf.fr/tomotools/nabu).

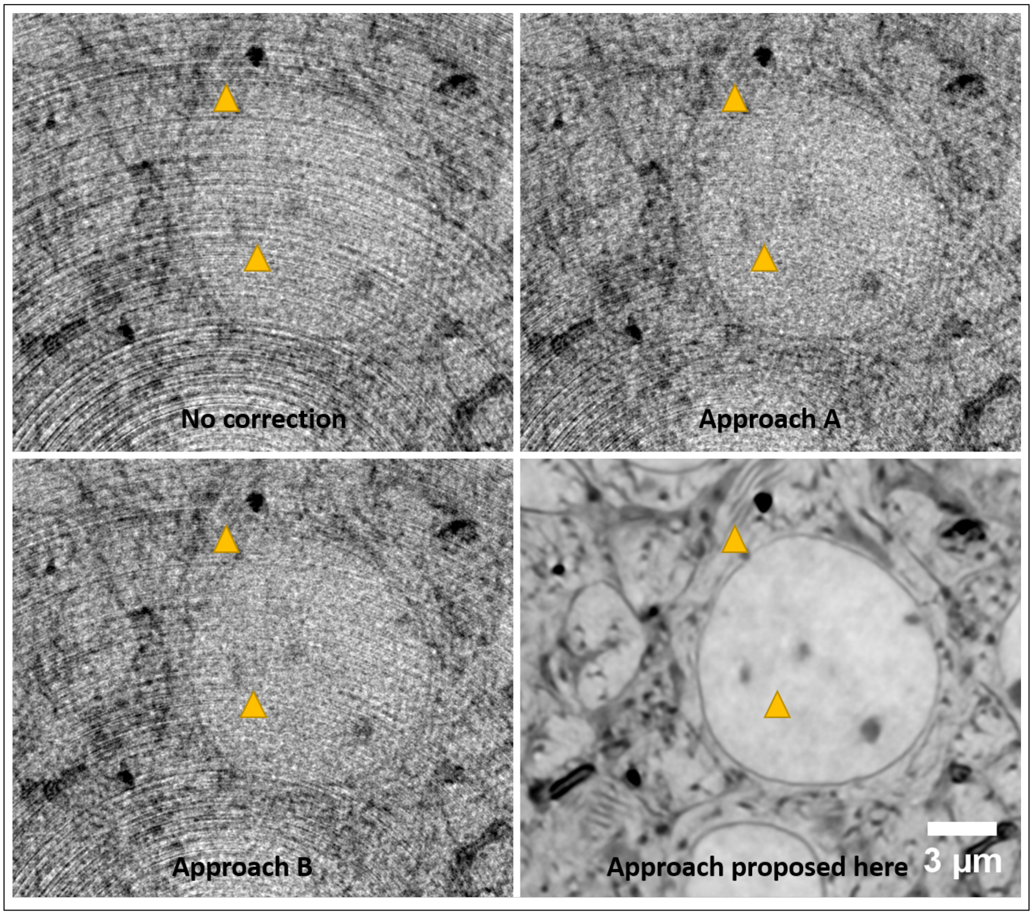


Fig. 5. Detailed views from the supplementary Figure **??**. Arrows point to endoplasmic reticulum and to one of the ring artifacts.

#### 7. Resolution improvement as function of training time

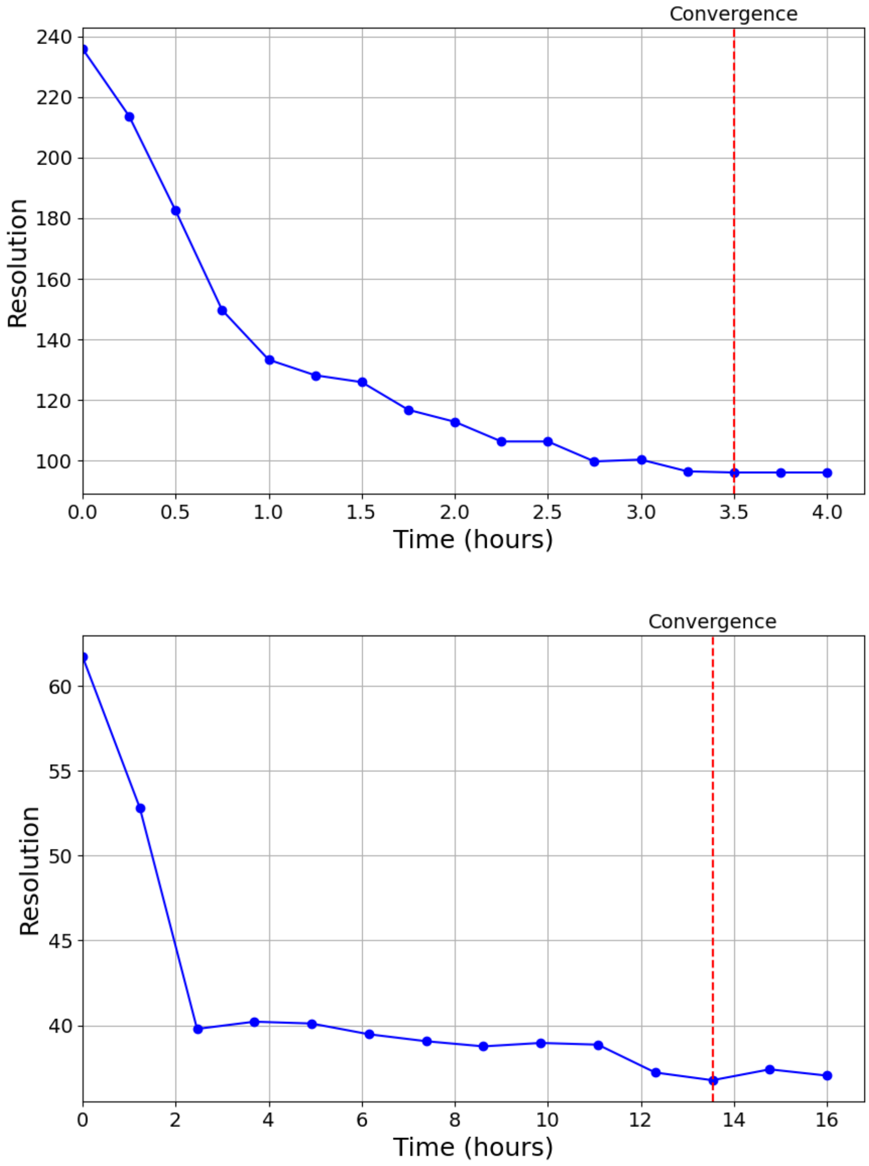


Fig. 6. Spatial resolution measured at different stages of training for the FSS (top) and OB15 (bottom) samples.

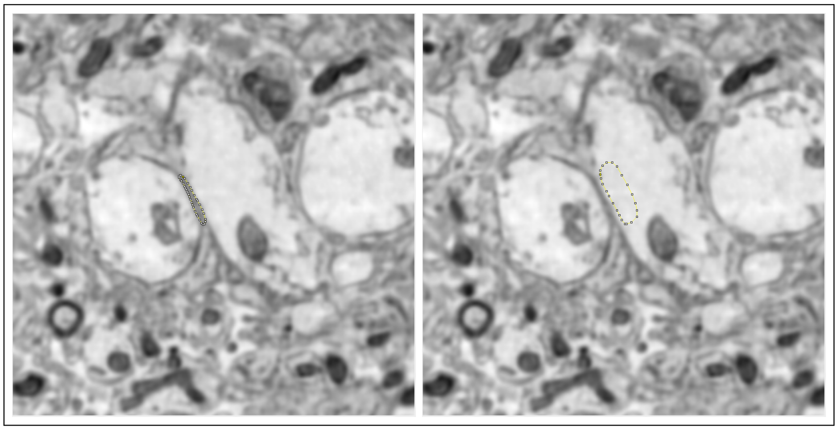

Fig. 7. Selection of image regions for computing CNR. Multiple regions from dendritic membranes (left panel) and cytosol (right panel) were manually selected to obtain the contrast to noise ratio measurements.

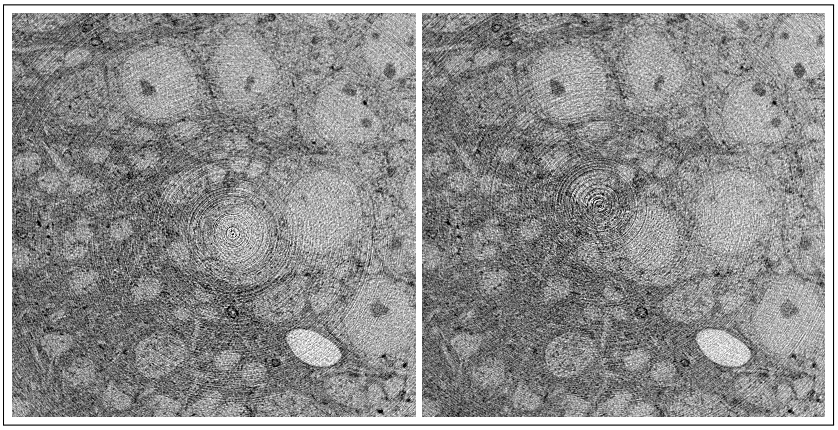

Fig. 8. Slices extracted from the training volumes used for ring correction showing the shifted location of the ring artifacts.

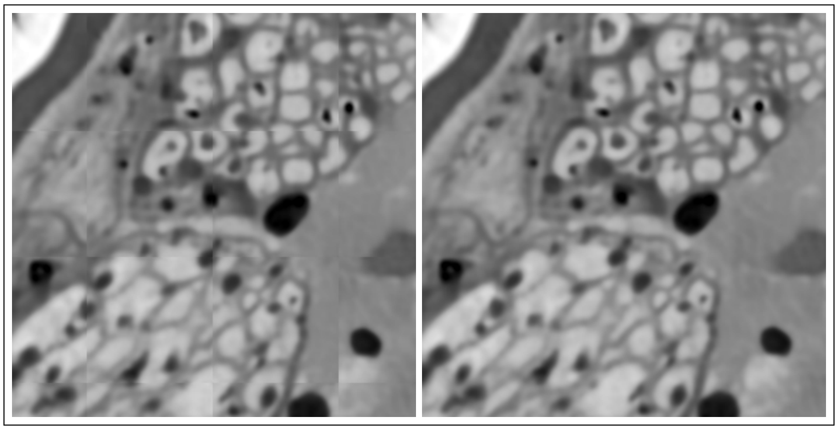

Fig. 9. FSS volume crops. Left: denoised image using a stride size equal to zero. Right: denoised image using a stride size of 80% of the 3D U-Net patch size. Grid artifacts visible on the left are no longer present on the right.

#### 8. Data split along propagation distances

Table 5. Denoising performances obtained using the 3D U-Net implementation trained on volumes generated with the distance projection partitioning strategy.

| Volume | Measured Resolution | CNR | Training Time |
|---|---|---|---|
| Raw OB25-dist | 104.6 | 3.2 ± 1.1 | - |
| Denoised OB25-dist | 55.7 | 7.0 ± 1.6 | 13h |

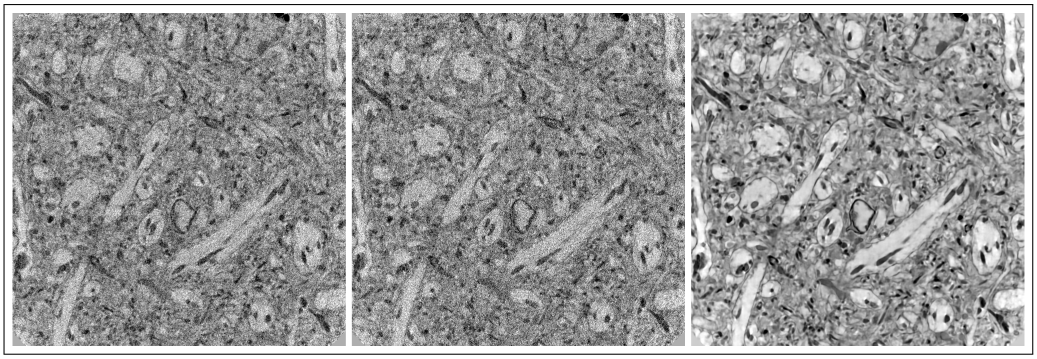

Fig. 10. Crops of the OB25 volume reconstructed using the distance projection partitioning strategy. Left and Middle: Raw training volumes. Right: denoised using the 3D U-Net implementation.
